# Culturing the ubiquitous freshwater actinobacterial acI lineage by supplying a biochemical ‘helper’ catalase

**DOI:** 10.1101/343640

**Authors:** Suhyun Kim, Ilnam Kang, Ji-Hui Seo, Jang-Cheon Cho

**Affiliations:** Department of Biological Sciences, Inha University, Incheon 22212, Republic of Korea

**Keywords:** acI lineage, Freshwater, Bacteria, Catalase, Hydrogen Peroxide, Cultivation

## Abstract

Unlike the ocean from which abundant microorganisms with streamlined genomes such as *Prochlorococcus*, *Pelagibacter*, and *Nitrosopumilus* have been isolated, no stable axenic bacterial cultures are available for the ubiquitous freshwater actinobacterial acI lineage. The acI lineage is among the most successful limnic bacterioplankton found on all continents, often representing more than half of all microbial cells in the lacustrine environment and constituting multiple ecotypes. Dilution-to-extinction culturing followed by whole-genome amplification recently yielded 20 complete acI genomes from lakes in Asia and Europe. However, stably growing pure cultures have not been established despite various efforts at cultivation using growth factors predicted from genome information. Here, we report two pure cultures of the acI lineage successfully maintained by supplementing the growth media with catalase. Catalase was critical for stabilizing growth by degrading hydrogen peroxide, irrespective of the genomic presence of the catalase-peroxidase (*katG*) gene, making the acI strains the first example of the Black Queen hypothesis reported for freshwater bacteria. The two strains, representing two novel species, displayed differential phenotypes and distinct preferences for reduced sulfurs and carbohydrates, some of which were difficult to predict based on genomic information. Our results suggest that culture of previously uncultured freshwater bacteria can be facilitated by a simple catalase-supplement method and indicate that genome-based metabolic prediction can be complemented by physiological analyses.

---

The acI lineage of the phylum *Actinobacteria* is the most abundant and cosmopolitan bacterial group in most freshwater environments. Since the acI lineage was first suggested to denote an abundant monophyletic actinobacterial group exclusively found in freshwater environments (1–3), many studies have demonstrated the ubiquity and prevalence of the acI lineage in diverse freshwater ecosystems on all continents (4–10).

Studies employing fluorescence *in situ* hybridization (FISH) and PCR-based 16S rRNA gene sequence profiling showed that the acI lineage and its subgroups exhibit specific distributions depending on the season, depth, and habitat characteristics and that there are >10 monophyletic tribes belonging to three sublineages (acI-A, -B, and -C) (4, 11–13). Ecophysiological studies using FISH combined with microautoradiography (MAR-FISH) showed that the acI lineage uptakes diverse substrates including leucine, thymidine, glucose, acetate, *N*-acetylglucosamine, and amino acid mixtures, but the substrate utilization patterns vary depending on the substrates, habitats, and sublineages (14–17). Shotgun metagenomics of freshwater samples (18, 19) and PCR-based surveys of enrichment cultures and single-amplified genomes (SAGs) of freshwater origin (20, 21) suggested that members of the acI lineage have genomes with low G+C content and carry genes for actinorhodopsin. Studies of SAGs (22, 23) and metagenome-assembled genomes (MAGs) (24) confirmed that the acI lineage genomes have low G+C content and encode actinorhodopsin genes and further suggested that the genomes are small-sized and enriched in genes for acquiring and utilizing carbohydrate and N-rich organic compounds.

As genomic data of the acI lineage has accumulated through culture-independent studies, cultivation of the acI lineage has become increasingly necessary because pure cultures can enable experimental verification of hypotheses deduced from the genomic data and reveal ecophysiological traits relevant to their survival and niche partitioning in natural habitats in a definitive manner (25, 26). The first report on cultivation of the acI lineage was based on a mixed culture obtained from a freshwater lake, but the proportion of acI cells was <6% (27).

Several enrichment cultures containing acI-B strains with higher proportions (up to >50%) were obtained by dilution culturing from freshwater lakes, and MAGs retrieved from these enrichment cultures suggested that the acI lineage depends on co-occurring microorganisms for the supply of various vitamins, amino acids, and reduced sulfur (28–30). Recently, Kang et al. (31) and Neuenschwander et al. (32) obtained transient axenic cultures of 20 acI strains in the acI-A, -B, and -C sublineages by dilution-to-extinction culturing. The complete genomes of these acI strains, obtained by whole-genome-amplification of the initial cultures, showed several characteristics of genome streamlining (33) such as small sizes (1.2–1.6 Mb), low G+C content, and high coding density. Metabolic reconstruction based on the genome sequences predicted auxotrophy of all strains for several vitamin B compounds and reduced sulfurs, as well as tribe-specific auxotrophy for some amino acids.

Although rich genomic data were obtained from putatively axenic cultures of the acI lineage after various cultivation efforts, all initial cultures failed to become stably-growing pure cultures (31, 32). This lack of pure cultures prevents further investigations of the physiology and ecology of this lineage, which dominates freshwater bacterioplankton, in contrast to marine environments where stably growing pure cultures of several ubiquitous prokaryotic groups including *Prochlorococcus* (34), *Pelagibacter* (35), and *Nitrosopumilus* (36) are available.

In this study, we report the first establishment of stably growing pure cultures of the acI lineage. Two acI strains belonging to different tribes were successfully cultured and maintained by lowering the concentration of hydrogen peroxide (H_2_O_2_) in the culture media by catalase supplementation, rendering the acI lineage the first case supporting the Black Queen hypothesis (BQH) (37) for freshwater heterotrophs. Using these novel pure cultures, we analyzed the phenotypic characteristics of the acI strains and determined metabolic features that are difficult to predict based only on genome information as well as features conforming to previous genome-guided inferences. Several physiological features differed between the two acI strains, providing insight into the niche separation and spatiotemporal dynamics of diverse acI populations.

## Results and Discussion

### Isolation of acI strains and efforts towards culture maintenance

Strain IMCC25003 of the acI-A1 tribe and strain IMCC26103 of the acI-A4 tribe (3, 4) were isolated from Lake Soyang in Korea by dilution-to-extinction culturing (31). Strains IMCC25003 and IMCC26103 were most closely related to the recently proposed Candidatus species ‘*Ca*. Planktophila sulfonica’ MMS-IA-56 (100%) and ‘*Ca*. Planktophila lacus’ MMS-21-148 (99.9%) (32), respectively, based on 16S rRNA gene phylogenetic analyses (*SI Appendix*, Fig. S1). According to the average nucleotide identity (ANI) (38) with their closest relatives (74–86%) (32), the two strains each represent novel species of the genus ‘*Ca*. Planktophila’.

Because the two strains are among the few acI isolates and the ability to maintain stably growing acI strains is prerequisite to studying the phenotypic characteristics of this lineage, we first attempted to resuscitate both strains from glycerol stocks using the same media used for isolation. IMCC26103 did not grow for 21 days after revival from the original frozen glycerol stocks but showed an abrupt increase in cell density (*SI Appendix*, Fig. S2*A*) because of the growth of contaminating strains (*SI Appendix*, Fig. S2*B*). IMCC25003 began growing without a lag from the original frozen stocks and reached a cell density of 7.6 × 10^5^ cells mL^-1^ in 5 days (*SI Appendix*, Fig. S2*C*; upper panel), but showed an atypical growth curve when the revived culture was transferred into the same medium on day 12 (*SI Appendix*, Fig. S2*C*; middle panel). When the first-transferred culture was inoculated again on day 15, no growth was detected until ~70 days (*SI Appendix*, Fig. S2*C*; lower panel).

Because maintenance of the growing cultures of acI strains was not successful in the media used for their original isolation, further attempts to obtain stable cultures were made by introducing various modifications to the media composition. These attempts were possible only for IMCC25003, as many glycerol stocks were stored using the initially revived cultures of IMCC25003. Modifications to media composition were based on the putative metabolic features and growth requirements of the acI lineage predicted by approaches such as MAR-FISH and single-cell genomics (15, 23). Culture media contained various supplements of FAMV medium (*see SI Appendix*, Table S1 for media composition) such as carbon substrates, proteinogenic amino acids, peptone, and yeast extract and different basal media including artificial freshwater medium (39) and spent medium of *Limnohabitans* sp. IMCC26003 (*SI Appendix*, Table S2). However, growth of IMCC25003 was not detected under any culture conditions tested.

### Successful maintenance of stably growing acI cultures by catalase addition

The failure to establish stable acI bacterial cultures by supplementation with defined or undefined nutrients led us to hypothesize that controlling the putative growth inhibitors may be more crucial than supplying growth promoters. Because the growth of *Prochlorococcus* and ammonia-oxidizing archaea was promoted when H_2_O_2_ was removed (40–42), we examined whether catalase, an enzyme that degrades H_2_O_2_, could facilitate the cellular growth of IMCC25003, although the IMCC25003 genome contains *katG*, which codes for a bifunctional catalase-peroxidase. IMCC25003 showed good growth and reached a maximum cell density of 2.7 × 10^7^ cells mL^-1^ in catalase (10 U mL^-1^)-amended medium (FAMV+CM+AA, *see* Materials and Methods for media composition), while no growth was detected in the medium devoid of catalase (*SI Appendix*, Fig. S3*A*). IMCC26103, which does not contain *katG*, was also successfully revived from the last-remained glycerol stocks in catalase-amended medium, reaching a maximum cell density of 5.2 × 10^7^ cells mL^-1^ (*SI Appendix*, Fig. S3*B*). Because both strains exhibited robust and reproducible growth upon repeated transfer into media supplemented with catalase, catalase was considered as a critical supplement in acI strain cultivation.

Before further experiments, dilution-to-extinction culturing of the successfully revived strains was performed twice to ensure the purity of the cultures, resulting in the establishment of stably growing pure cultures. The purity of the established cultures was verified by FISH (Fig. 1*A*), transmission electron microscopy (Fig. 1*B*), and whole genome sequencing followed by read mapping. Both strains produced reddish cell pellets because of the presence of carotenoids and actinorhodopsin in 4-L cultures (*SI Appendix*, Fig. S4*A*), which were subsequently used for genome sequencing. The genome sequences were identical (1 base pair difference) to those obtained by multiple displacement amplification in our previous study (31), with more even sequencing coverages (*SI Appendix*, Fig. S4*B*). For both strains, more than 99.9% of sequencing reads were mapped to the complete genomes, demonstrating that the cultures were pure.

**Fig. 1.**
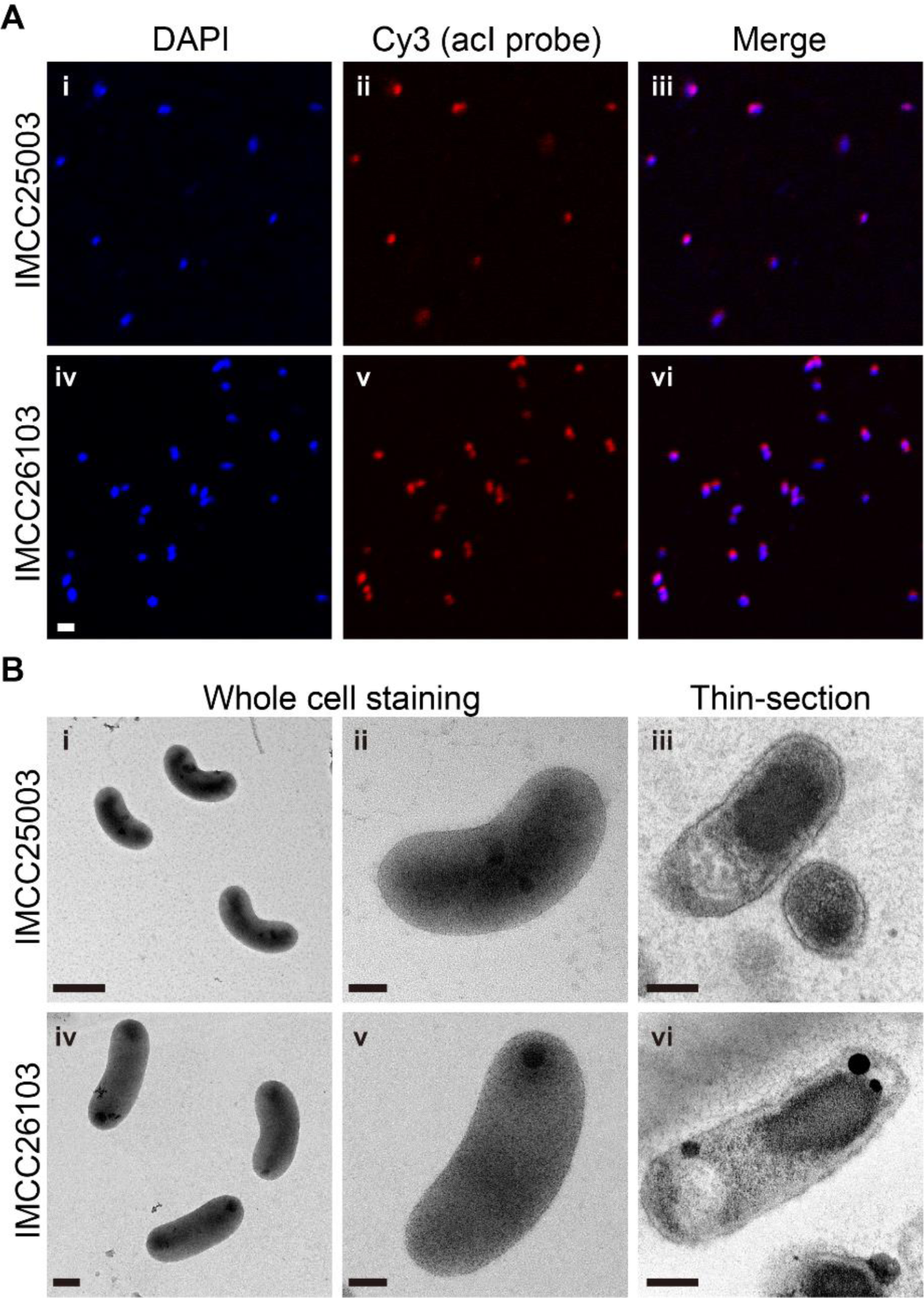
Microscopic examination of the two acI strains for purity evaluation and morphological characterization. (*A*) Demonstration of culture purity of strains IMCC25003 (upper; i, ii, and iii) and IMCC26103 (lower; iv, v, and vi) by FISH analysis. The acI cells were dual-stained with DAPI (i and iv) and Cy3-conjugated oligonucleotide probes specific to the acI lineage (AcI-852 and AcI-1214) (ii and v). Merged images are shown in iii and vi. Bar, 1 μm. (*B*) Cell morphology of strains IMCC25003 (upper; i, ii, and iii) and IMCC26103 (lower; iv, v, and vi) examined by transmission electron microscopy. The acI cells were stained with uranyl acetate (i, ii, iv, and v) or double stained with uranyl acetate and lead oxide after thin-sectioning (iii and vi). Scale bars; 0.5 μm (i), 0.2 μm (iv), and 0.1 μm (ii, iii, v, and vi).

To reconfirm the catalase-dependent growth, the two acI strains were cultivated with various concentrations of catalase (0–20 U mL^-1^). Growth of the strains was dependent on the presence of catalase, as no growth was detected in the absence of catalase, while 0.5 U mL^-1^ catalase increased the cell density to ~10^7^ cells mL^-1^ (Fig. 2 *A* and *B*). Cell densities and H_2_O_2_ concentrations were simultaneously monitored in a catalase-spiked growth experiment to determine whether the pivotal role of catalase in promoting growth was mediated by degradation of H_2_O_2_. Without catalase, no cellular growth was detected (Fig. 2 *C* and *D*) and the H_2_O_2_ concentration was maintained at the initial level of ~60 nM (Fig. 2 *E* and *F*). In contrast, when 10 U mL^-1^ catalase was added at the time of inoculation, the H_2_O_2_ concentration rapidly decreased to less than 10 nM and the acI strains began growing, reaching maximum cell densities in 6–9 days. When catalase was spiked at 7 or 13 days for IMCC25003 and 5 or 12 days for IMCC26103, the H_2_O_2_ concentration rapidly decreased from ~60 to ~10 nM (Fig. 2 *E* and *F*). The acI strains, which showed no growth before catalase addition, began growing following the decrease in H_2_O_2_ concentration immediately after catalase addition, indicating that H_2_O_2_ removal by catalase was critical for maintaining the growth of the acI strains.

**Fig. 2.**
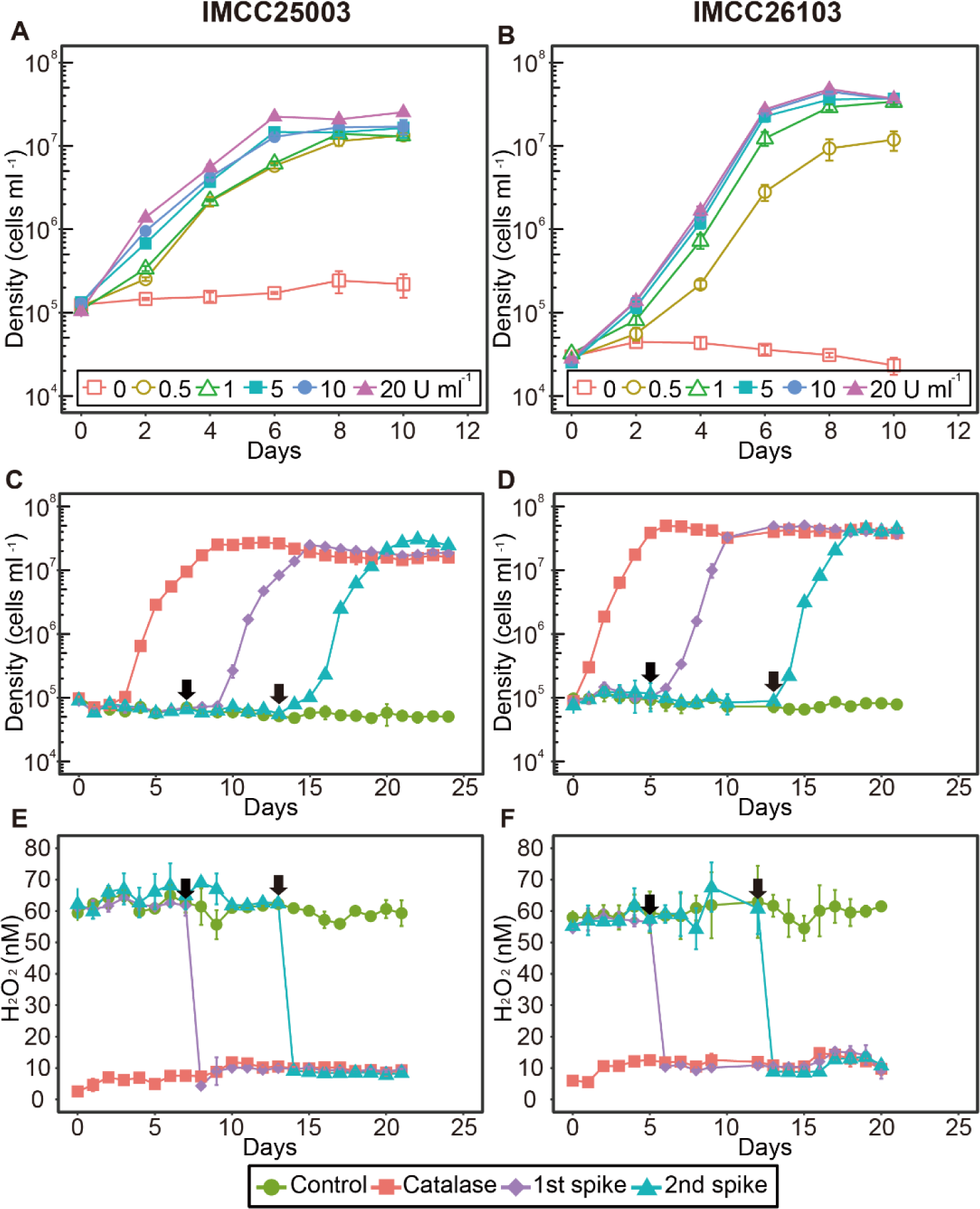
Catalase-dependent growth of the two acI strains. (*A* and *B*) The effect of various concentrations (0–20 U mL^-1^) of catalase on the growth of strains IMCC25003 (*A*) and IMCC26103 (*B*). (*C*–*F*) The effect of catalase addition on cell densities (*C* and *D*) and H_2_O_2_ concentrations (*E* and *F*) during cultivation of strains IMCC25003 (*C* and *E*) and IMCC26103 (*D* and *F*). Catalase (10 U mL^-1^) was added to the culture media at day 0 (‘Catalase’) or days indicated by black arrows (‘1st spike’ and ‘2nd spike’). No catalase was added to the cultures designated as ‘Control’. All experiments were performed in triplicate using the culture medium FAMV+CM+AA (Table S1). Error bars indicate standard deviation. Note that error bars shorter than the size of the symbols are hidden.

Among the several traits explained by the BQH (37) such as auxotrophy for vitamins or amino acids (33, 43), the dependency on external H_2_O_2_ scavengers is the first-reported and among the most representative cases of the BQH, discovered in two abundant marine autotrophic prokaryotic groups, *Prochlorococcus* and ammonia-oxidizing archaea (40–42). The dependency of cellular growth on the external H_2_O_2_ scavenger found in this study makes the acI lineage the first freshwater bacterial example supporting the BQH, which explains the evolution of metabolic dependency of genome-streamlined oligotrophic bacteria on external ‘helpers’ in nutrient-depleted environments (33, 44). This crucial role of catalase in cultivating the acI strains was unexpected because all culture media used in cultivation trials contained pyruvate, a well-known chemical scavenger of H_2_O_2_ (45, 46) and because acI bacteria failed to resuscitate from the original dilution cultures regardless of the presence of *katG* (31, 32). In this context, we first assumed that unknown growth factors were required for laboratory cultivation rather than H_2_O_2_ scavengers and thus we tested various nutrients and growth factors to revive the acI strains. However, only catalase treatment resulted in successful cultivation of the acI strains, while all other attempts failed. Our cultivation results suggest that acI bacteria grow only at very low concentrations of H_2_O_2_ attainable by catalase addition under laboratory conditions, regardless of the presence of *katG*, suggesting that the metabolic dependency of bacterial groups cannot be predicted based solely on genome information.

### Characterization of KatG of the acI lineage

Although *katG* was found only in IMCC25003, the two acI strains showed similar growth responses to H_2_O_2_, with no growth at high concentrations of H_2_O_2_ but good growth at minute concentrations of H_2_O_2_ lowered by catalase. This result indicates that *katG* in IMCC25003 or its gene product, bifunctional catalase-peroxidase (KatG), did not function properly to defend against H_2_O_2_ in the culture experiments. Therefore, we first examined the expression of *katG* in IMCC25003 by qPCR. The qPCR results showed that *katG* was expressed and the relative expression level increased slightly with increasing H_2_O_2_ concentrations (*SI Appendix*, Fig. S5). Next, *katG* of IMCC25003 was cloned and expressed in *E. coli* and the gene product was purified by affinity chromatography (*SI Appendix*, Fig. S6). The native molecular weight of IMCC25003 KatG was ~165 kDa (*SI Appendix*, Fig. S6*D*) and molecular weight of the subunit analyzed by SDS-PAGE was ~83 kDa (*SI Appendix*, Fig. S6*A*), indicating that IMCC25003 KatG formed a homodimer composed of two subunits. In an in-gel enzyme activity assay, IMCC25003 KatG showed both catalase and peroxidase activities but its catalase activity was much lower than that of bovine catalase (KatE), which belongs to the monofunctional catalase group and was supplemented into the culture media in this study (*SI Appendix*, Fig. S7). To more accurately assay the catalase activity of IMCC25003 KatG, decomposition of H_2_O_2_ was plotted using varying amounts of enzymes (*SI Appendix*, Fig. S8). All enzymatic parameters including specific activity (U mg^-1^) and catalytic efficiency (*k_cat_*/*K_m_*) showed that IMCC25003 KatG displayed lower catalase activities than those of other microbial KatGs reported previously, as well as KatE (*SI Appendix*, Table S3), suggesting that the activity of IMCC25003 KatG is not sufficient to overcome the growth inhibition by H_2_O_2_ present in the culture media.

We investigated the distribution of *katG* among available acI genomes, as only one (IMCC25003) of four acI strains isolated from Lake Soyang contains *katG* (31). *katG* was detected in 14 of 20 complete acI genomes (*SI Appendix*, Table S4). When single-amplified genomes were included, 20 of 35 genomes carried *katG*. The distribution of *katG* in the acI lineage was biased toward acI-A compared to acI-B. Of the 25 acI-A genomes, 19 carried *katG*s, while only one of 10 acI-B genomes contained *katG* (*SI Appendix*, Table S4); thus, we analyzed the phylogenetic features of acI KatG proteins. Unexpectedly, phylogenetic analyses including diverse bacterial KatG proteins (47) indicated that KatGs of the acI lineage belonged to two distinct groups that are separated widely in the entire KatG tree (Fig. 3*A*). Further, this KatG phylogeny of the acI lineage was not consistent with a phylogenomic tree of the lineage based on conserved proteins (Fig. 3*B*). KatGs of the acI-A sublineage were divided into two different KatG groups depending on the tribes. acI-A1 and A2 KatGs formed group A, while acI-A4, A5, A6, and A7 KatGs formed group B together with KatG of acI-B4 (Fig. 3). These results indicate that complicated evolutionary processes occurred for KatG in the acI lineage. Because KatG of IMCC25003 belonging to the group A showed low activity, it would be interesting to determine if KatGs belonging to group B also exhibit low catalase activity, as isolates containing group B KatG did not grow in lake water-based medium (32).

**Fig. 3.**
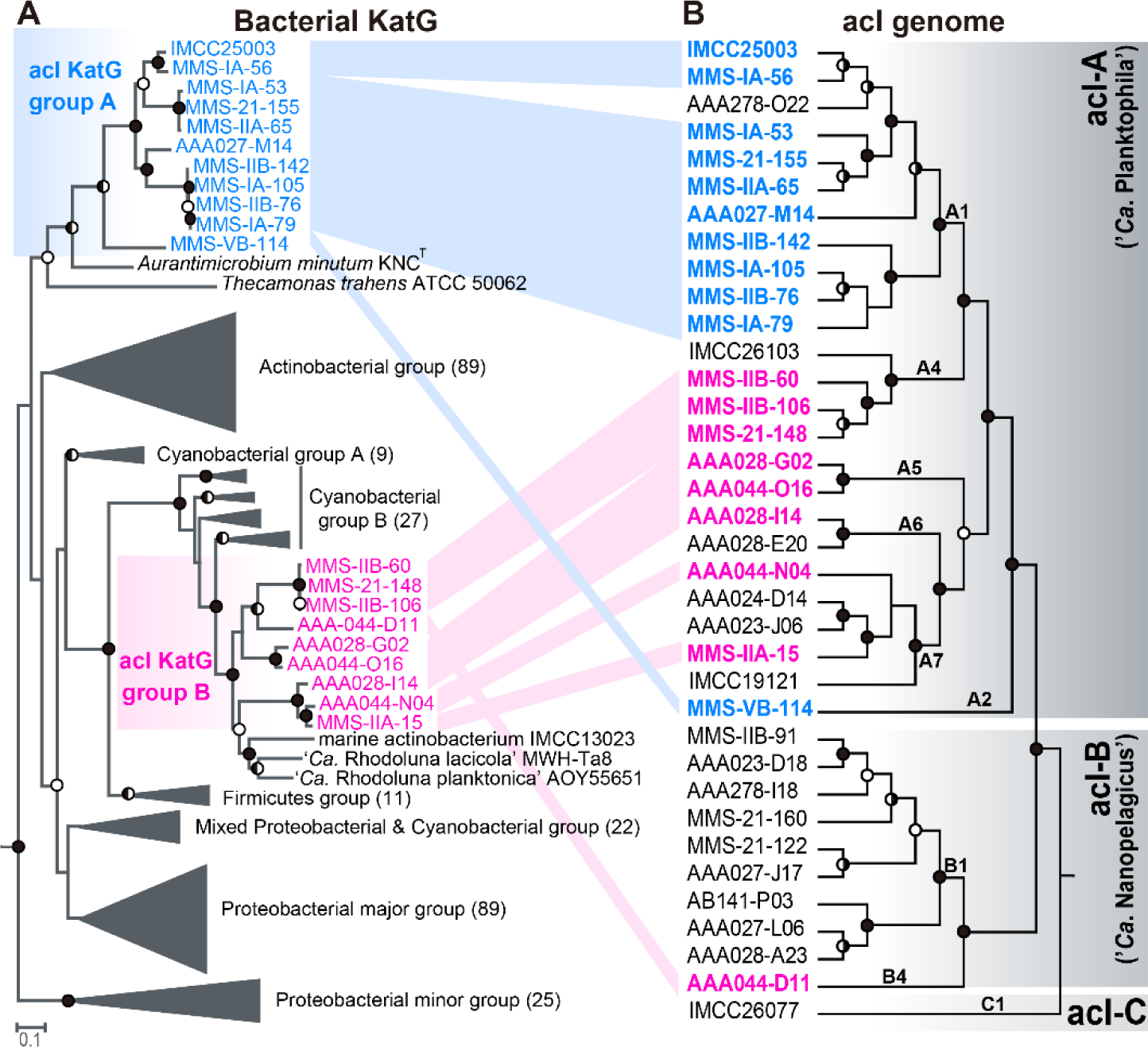
Comparison between a phylogenetic tree of bacterial KatG proteins and phylogenomic tree of members of the acI lineage. (*A*) A maximum-likelihood tree of bacterial KatG protein sequences including those predicted in the acI genomes. The acI KatG proteins are indicated in color (blue and red for acI KatG group A and B, respectively), while other bacterial KatG proteins were grouped following the scheme proposed by Zamocky *et al.* (47). The number of proteins included in each KatG group are indicated within parentheses at the end of group names. The KatG sequences of *Rhodopirellula baltica* and *Pirellula staleyi* (both from the phylum *Planctomycetes*) were used as the outgroup. (*B*) A phylogenomic tree of members of the acI lineage constructed using concatenated alignment of conserved marker proteins. The acI strains containing *katG* are indicated in color according to the grouping in the left KatG tree. The SAG AAA027-D23 (belonging to the acSTL-A1 tribe) (30) was set as an outgroup. Only the tree topologies are shown, and branch lengths do not represent phylogenetic distances. For both trees, bootstrap values (from 100 and 250 replicates for the KatG tree and phylogenomic tree, respectively) are shown at the nodes as filled circles (≥90%), half-filled circles (≥70%), and empty circles (≥50%).

### Phenotypic characterization of the acI strains

Establishment of stable pure cultures by catalase addition enabled the first phenotypic characterization of the acI lineage, including detailed morphology (Fig. 1*B*), temperature preference (*SI Appendix*, Fig. S9), fatty acid profile (*SI Appendix*, Table S5), growth requirement, and substrate utilization (Fig. 4), leading to the proposal of two novel species named as ‘*Ca*. Planktophila rubra’ (ru’bra. L. fem. adj. *rubra* reddish, pertaining to the reddish color of cells) for IMCC25003 and ‘*Ca*. Planktophila aquatilis’ (a.qua.ti′lis. L. fem. adj. *aquatilis* living, growing, or found, in or near water, aquatic) for IMCC26103 (*see SI Appendix*, *Supplementary Methods*). Actively growing cells of the two strains were curved rods of very small sizes and biovolumes of 0.041 (for IMCC25003) and 0.061 µm^3^ (for IMCC26103), representing some of the smallest ultramicrobial (<0.1 µm^3^) cells among known cultivated freshwater bacteria, but 1.6–2.4-fold larger cell volumes than those of other acI bacteria estimated by epifluorescence microscopy (32) (*SI Appendix*, Table S6).

**Fig. 4.**
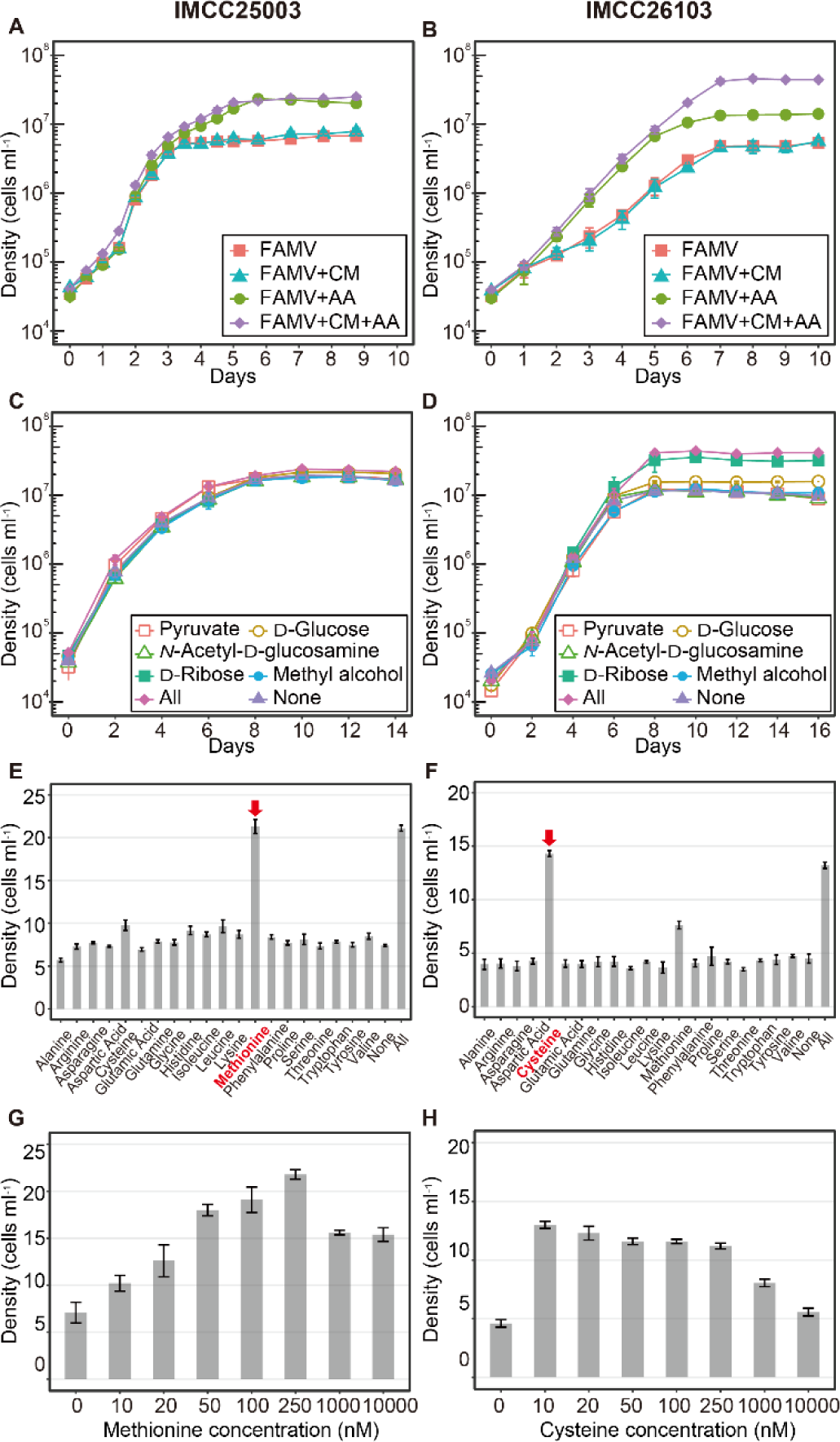
Effects of carbon sources and amino acids on the growth of the acI strains IMCC25003 (figures on the left: *A*, *C*, *E*, and *G*) and IMCC26103 (figures on the right: *B*, *D*, *F*, and *H*). (*A* and *B*) Effects of amino acid mixture (AA) and/or carbon source mixture (CM). AA and CM were added to the FAMV medium separately or together. Detailed medium composition is presented in Table S1. (*C* and *D*) Effects of individual carbon compound. Each of 5 carbon compounds [5 μM, except for pyruvate (50 μM)] was added to the FAMV+AA medium. All, 5 carbon substrates were added together; None, no carbon source was added. (*E* and *F*) Effects of individual amino acids. Each of 20 proteinogenic amino acids was added at a concentration of 100 nM to the basal medium FAMV. All, 20 amino acids were added together; None, no amino acid was added. (*G* and *H*) Optimal concentrations of L-methionine (for IMCC25003) and L-cysteine (for IMCC26103). Various concentrations (0–10,000 nM) of L-methionine or L-cysteine were added to the basal medium FAMV.

The cell densities of IMCC25003 were increased by amino acids but not by carbon mixtures (Fig. 4*A*). Growth of IMCC26103 was enhanced by both amino acids and carbon mixtures, but the enhancement by carbon mixtures was observed only in the presence of amino acids (Fig. 4*B*). These results indicate that the growth of both strains was mainly limited by at least one of 20 amino acids and that IMCC26103 (but not IMCC25003) utilized at least one of the carbon sources added. In each carbon compound-amended experiment, no single carbon compound affected the growth of IMCC25003 (Fig. 4*C*), while ribose and glucose enhanced the growth of IMCC26103 (Fig. 4*D*). These results were supported by genomic information showing that the IMCC26103 genome contained genes for D-ribose pyranase, ribokinase, and glucose ABC transporter, while these genes were not found in the IMCC25003 genome. In each amino acid-amended test, methionine and cysteine increased the growth of both strains (Fig. 4 *E* and *F*), supporting the genome-based metabolic prediction that acI strains lack an assimilatory sulfate reduction pathway and thus require reduced sulfur compounds for growth (31).

Interestingly, the two strains showed different preferences for reduced sulfur compounds. IMCC25003 preferred methionine (optimum at 250 nM) over cysteine (Fig. 4 *E* and *G*), while IMCC26103 preferred cysteine (optimum at 10 nM) over methionine (Fig. 4 *F* and *H*), which was difficult to predict from genome information. Although a limited number of substrates was tested, these differential preferences of different acI tribes on growth substrates may lead to niche differentiation, underlying the tribe-specific spatiotemporal dynamics of the acI lineage found in several cultivation-independent studies (6, 30, 48, 49).

### Concluding remarks

We successfully maintained actively-growing pure cultures of the acI lineage, the most abundant freshwater bacterioplankton group, by supplementing catalase into the culture media to lower H_2_O_2_ concentration, enabling further analysis of the physiological properties that could not be inferred from genome sequences alone. This simple catalase-supplement method may accelerate the cultivation of bacterioplankton with streamlined genomes and thus contribute to studies of the ecological roles of ubiquitous and abundant freshwater oligotrophic bacteria.

## Materials and Methods

Please see *SI Appendix* for experimental details on “Measurement of bacterial cell densities”, “Measurement of *katG* expression by qPCR”, “Expression, purification, and characterization of KatG from IMCC25003”, and “Phylogenetic analyses based on 16S rRNA gene, whole genome, and KatG protein”.

### Initial isolation of the acI stains, medium preparation, and revival experiments

Freshwater collection, initial isolation, and identification of strains IMCC25003 and IMCC26103 were described in our previous study (31). Briefly, both strains were initially isolated as liquid cultures by high-throughput culturing method based on dilution-to-extinction in 0.2 μm-filtered and autoclaved freshwater media supplemented with very low levels of carbon compounds, amino acids mixture, and vitamin mixture. Two cryovials each containing 200 μL of initial high-throughput cultures suspended in 10% (v/v) sterile glycerol were stored at −80°C and used for further revival experiments. As shown in *SI Appendix*, Fig. S2, the axenic culture of IMCC25003 was obtained from the revival experiments, which was further used in the following experiments to establish stably maintained pure cultures of IMCC25003.

Culture media for reviving IMCC25003 were prepared based on natural freshwater (*see SI Appendix*, Table S1 for culture media ingredients). Natural freshwater collected at a depth of 1 m from the Dam station (37°56ʹ50.6″ N, 127°49ʹ7.9″ E) of Lake Soyang in February 2016 was filtered through a 0.2-μm pore-size membrane filter (Supor, Pall Corporation), autoclaved for 1.5 h, and aerated for 3 h. The filtered-autoclaved-aerated medium was supplemented with 10 μM NH_4_Cl, 10 μM KH_2_PO_4_, and trace metal mixture (50), which was designated as FAM (filtered and autoclaved freshwater medium). The FAM medium supplemented with the vitamin mixture (51) (*SI Appendix*, Table S1), designated as FAMV, was used as a basal medium that did not contain extra-amended organic compounds. FAMV supplemented with a carbon mixture (CM; 50 μM pyruvate, 5 μM D-glucose, 5 μM *N*-acetyl-D-glucosamine, 5 μM D-ribose, and 5 μM methyl alcohol) and amino acid mixture (AA; 20 proteinogenic amino acids, 100 nM each) were prepared and used as basal heterotrophic growth media.

As the first attempt to obtain a stable culture of IMCC25003 (*SI Appendix*, Table S2), a 100-μL glycerol stock of IMCC25003 was inoculated into 20 mL of FAMV, FAMV amended with CM (FAMV+CM), FAMV amended with AA (FAMV+AA), and FAMV amended with CM and AA (FAMV+CM+AA). Different concentrations (0.5×, 1×, 5×, and 10×) of CM were added to FAMV+AA and the growth of IMCC25003 was tested. A 0.1-μm-filtered but non-autoclaved freshwater medium (FM) and artificial freshwater medium (39) each supplemented with vitamin mixture, CM, and AA were also tested. In the second attempt, the following organic nutrients were individually supplemented into FAMV+CM+AA: 20 μM acetate, 20 μM oxaloacetate, 20 μM putrescine, 20 μM glycerol, 20 μM xylose, 1 mg L^-1^ proteose peptone, and 1 mg L^-1^ yeast extract. In the third attempt, the spent medium of the genus *Limnohabitans* which is often considered as a co-occurring bacterium with acI bacteria, was added to the growth test medium. *Limnohabitans* sp. IMCC26003 grew to 3.1 × 10^6^ cells mL^-1^ in 100 mL of FAMV+CM+AA at 25°C for 2 weeks. After the *Limnohabitans* culture was filtered through a 0.2-μm pore-size membrane followed by a 0.1-μm pore-size membrane, 1 mL of the filtrate of the spent medium and 100 μL of glycerol stock of IMCC25003 were inoculated into 20 mL of FAMV+CM+AA. In the fourth attempt, catalase (from bovine liver, C9322, Sigma-Aldrich) stock solution (10^5^ U mL^-1^ in 10 mM PBS, pH 7.4) was added to FAMV+CM+AA at a final concentration of 10 U mL^-1^ and 100 μL glycerol stock of IMCC25003 was inoculated. All reviving tests were performed at 18°C for 5 weeks.

### Culture maintenance and confirmation of culture purity

Bacterial cultures of strains IMCC25003 and IMCC26103 revived first from glycerol stocks by catalase amendment were purified twice by dilution-to-extinction culturing using FAMV+CM+AA containing 10 U mL^-1^ of catalase. Cells were diluted to 5, 1, or 0.1 cells mL^-1^ and dispensed into 48-well plates (1 mL per each well). After incubation, growth-positive wells (>10^5^ cells mL^-1^) from the most diluted inoculum (0.1 cells mL^-1^) were used for next round of dilution-to-extinction, resulting in the establishment of stably-growing pure cultures.

To confirm the purity of the established pure cultures by FISH, 5 mL of exponentially grown cells were fixed with 2% paraformaldehyde (in PBS, pH 7.4) and filtered through 0.2-μm polycarbonate filters (Isopore, Millipore). Hybridization was performed at 35°C for 6 h in hybridization buffer [900 mM NaCl, 20 mM Tris (pH 7.4), 0.01% SDS, 15% formamide] with Cy3-labeled oligonucleotide probes (AcI-852 and AcI-1214; 2 ng μL^-1^ each) and helper probes (AcI-852-H1 and AcI-852-H2; 2 ng μL^-1^ each) targeting the acI lineage (52). After washing the membranes twice with washing buffer [150 mM NaCl, 20 mM Tris (pH 7.4), 6 mM EDTA, and 0.01% SDS] at 55°C for 10 min followed by DAPI (4,6-diamidino-2-phenylindole) staining, Cy3-positive and DAPI-positive cells were visualized with a confocal laser scanning microscope (LSM 510 META, Carl Zeiss).

For purity confirmation by genome sequencing, the two strains were cultivated in 4 L of FAMV+CM+AA supplemented with 10 U mL^-1^ of catalase. Cells were harvested by centrifugation at 20,000 ×*g* for 120 min and the cell pellets (*SI Appendix*, Fig. S4*A*) were used for genomic DNA extraction. Sequencing libraries were constructed using the Nextera library preparation kit (Illumina). Genome sequencing was performed on an Illumina MiSeq platform (2 × 300 bp) by ChunLab, Inc (Seoul, Republic of Korea). Raw sequencing data were assembled using SPAdes 3.9.0 (53), resulting in a circularly closed complete genome for both strains.

Comparison of the genome sequences obtained in this study to those obtained by multiple displacement amplification (31) was performed using BLASTn. Calculation of sequencing coverages and estimation of culture purity based on reads mapping were performed as described by Kang *et al*. (31), using the ‘depth’ and ‘flagstat’ options in samtools.

After purification, both strains were stored as 10% (v/v) glycerol suspensions at −80°C and working cultures were maintained in FAMV+CM+AA supplemented with 1 U mL^-1^ of catalase at 25°C. Throughout this study, the peak shape of the cultures in the flow cytometry, morphology of SYBR Green I-stained cells in epifluorescence microscopy, FISH images obtained using acI-specific probes, and electropherograms from 16S rRNA gene sequencing were routinely examined to evaluate culture purity.

### Characterization of catalase-dependent growth properties

The effects of the catalase concentration on the growth of strains IMCC25003 and IMCC26103 were tested in triplicate in FAMV+CM+AA supplemented with 0.5, 1, 5, 10, and 20 U mL^-1^ of catalase at 25°C. Bacterial cell densities were measured by flow cytometry every 2 days.

To confirm that catalase enhanced cellular growth and decreased H_2_O_2_ concentration, 10 U mL^-1^ of catalase was added to IMCC25003-inoculated culture medium (FAMV+CM+AA) that had been maintained for 0, 7, and 13 days without catalase treatment. Similarly, for strain IMCC26103, 10 U mL^-1^ of catalase was spiked at 0, 5, and 12 days. Cultures maintained without catalase were used as controls. For all cultures, cell densities were monitored every day by flow cytometry. At the same time, the concentration of H_2_O_2_ in the culture medium was measured using the modified acridinium ester chemiluminescence method (54) described below.

### Determination of hydrogen peroxide concentration

A modified acridinium ester chemiluminescence method (54) was used to determine H_2_O_2_ concentrations in the culture medium. To measure H_2_O_2_ concentrations at the nanomolar level, acridinium NHS ester (A63063; Cayman Chemical) was used as an indicator and the resulting chemiluminescence was estimated using a SpectraMax L microplate Reader (Molecular Devices). A standard curve for H_2_O_2_ was prepared in the range of 0.02–2.0 μM by diluting 30% (v/v) of H_2_O_2_ solution (H1009; Sigma-Aldrich) with ddH_2_O. The acridinium ester-H_2_O_2_ response signal was integrated for 13 s after sequential injections of 20 μL 2 M Na_2_CO_3_ (pH 11.3) followed by 80 μL of 2.2 mg L^-1^ acridinium ester (in 1 mM phosphate buffer, pH 3) at 300 μL s^-1^ injection flow speed and 0.1-s interval. All measurements were conducted in triplicate at 20°C.

### Morphological and physiological characterization of acI strains

Cell morphology was examined by transmission electron microscopy (CM200, Philips) using whole cell staining and thin-sectioning. For whole cell staining, 20-mL cultures were gently filtered using a 0.2-μm pore size polycarbonate membrane on which formvar/carbon-coated copper grids were placed, followed by staining of the grids with 0.5% uranyl acetate. To prepare thin-sectioned samples, cell pellets were harvested by centrifugation of 2-L cultures at 20,000 ×*g* for 120 min. Cell pellets that were primary fixed with 2.0% Karnovsky’s fixative and secondary fixed with 1.0% osmium tetroxide were dehydrated using ethanol series from 35% to 100% and Epon 812 resin was infiltrated. Ultrathin sections (70-nm) were prepared with an ultramicrotome using a diamond knife, placed on formvar/carbon-coated copper grids, and double-stained with 2% uranyl acetate and 0.5% lead oxide.

The temperature range for growth of the two acI strains was monitored at 10–35°C in FAMV+CM+AA supplemented with 10 U mL^-1^ of catalase. After the optimum temperature had been determined, all growth experiments were performed in triplicate at 25°C in different growth media supplemented with 10 U mL^-1^ of catalase. The growth curves in FAMV, FAMV+CM, FAMV+AA, and FAMV+CM+AA were determined to evaluate the effects of carbon compound mixtures and amino acids mixtures on cellular growth. To identify which amino acids are required for growth, 20 individual amino acids were supplemented in the FAMV medium at 100 nM and cellular growth was monitored. Similarly, 5 individual carbon sources (50 μM pyruvate, 5 μM D-glucose, 5 μM *N*-acetyl-D-glucosamine, 5 μM D-ribose, and 5 μM methyl alcohol) were supplemented into FAMV+AA and the growth curves were determined. To determine the optimal concentration of reduced sulfur compounds, growth of the acI strains was measured at 0–10 μM of methionine (for IMCC25003) and cysteine (for IMCC26103).

Cellular fatty acid profiles were determined using cells harvested from 2-L cultures grown in FAMV+CM+AA supplemented with 10 U mL^-1^ of catalase. Fatty acid methyl esters were extracted from cell pellets using the standard protocol for the Sherlock Microbial Identification System (MIDI, Inc.) and analyzed by gas chromatography (Agilent 7890 GC) based on MIDI version 6.1 with the TSBA6 database (55).

### Data availability

The complete genome sequences of strains IMCC25003 and IMCC26103 obtained from cell pellets using the Illumina MiSeq platform have been deposited in GenBank with the accession numbers CP029557 for IMCC25003 and CP029558 for IMCC26103.

## ACKNOWLEDGEMENTS

This study was supported by the Mid-Career Research Program (to J-CC, No. NRF-2016R1A2B2015142) and Science Research Center grant (to J-CC, No. NRF-2018R1A5A1025077) through the National Research Foundation (NRF) funded by the Ministry of Sciences and ICT, and by the Basic Science Research Program funded by the Ministry of Education, Republic of Korea (to IK, No. NRF-2016R1A6A3A11934789; to J-HS, No. NRF-2016R1A6A3A11935361).

